# The eukaryome of modern microbialites reveals distinct colonization across aquatic ecosystems

**DOI:** 10.1101/2023.10.14.562355

**Authors:** Anthony Bonacolta, Pieter T Visscher, Javier del Campo, Richard Allen White

**Author notes:** Address correspondence to Richard Allen White III & Javier del Campo.

## Abstract

Microbial diversity includes bacteria, archaea, eukaryotes, and viruses; however, protists are less studied for their impact and diversity within ecosystems. Protists have been suggested to shape the emergence and decline of ancient stromatolites. Modern microbialites offer a unique proxy to study the deposition of carbonate by microbial communities due to analog status for ancient ecosystems and their cosmopolitan abundance. We examined protists across aquatic ecosystems between freshwater (Kelly and Pavilion Lake in British Columbia, Canada) and marine microbialites (Shark Bay, Australia and Highborne Cay, Bahamas) to decipher the transition with respect to diversity and composition. While factors such as sequencing technology and primer-bias might influence our conclusions, we found that at the taxonomic compositional-level, the freshwater microbialite communities were clearly distinct from the marine microbialite communities. Chlorophytes were significantly more abundant in the freshwater microbialites, while saltwater microbialites communities were primarily composed of pennate diatoms. Despite the differences in taxonomic make-up, we can infer the convergent important role of these protists to microbialite community health and function. These results highlight not only the consistency and potential role of microbialite eukaryotic communities across geographic locations, but also that other factors such as salinity seem to be the main drivers of community composition.

## Introduction

Microbialites represent the oldest known persistent ecosystems and potentially provide the earliest evidence of life [1, 2]. The geologic record indicates that microbialite diversification occurred concurrently with the emergence of life ∼3.50 billion years ago (Gya). This was followed by a period of dominance lasting until ∼542 million years ago (Mya), the period of the Cambrian explosion [1, 3–5]. Microbialites have, therefore, persisted for ∼85% of geological history [3, 6] Modern microbialites may grant insight into the oldest aspects of the biosphere [2, 3].

It has been generally thought that microbialites are rare environments within the biosphere. It has been recently shown that microbialites represent a cosmopolitan distribution across the globe [7]. Microbialites are found in water contains stable alkaline pH (>8), as free divalent cations (>7.5 gpg Ca^2+^ or Mg^2+^), is relatively hard water (≥ 121 mg CaCO_3_ L^-1^), oligotrophic, clear water with visible light penetration, and dissolved inorganic carbon (DIC) [8]. The variety of ecosystems include soil, caves, hot springs, lakes, oceans, hypersaline lagoons, tropical lagoons, and remnant mining sites [7, 9, 10].

Eukaryotes in microbialites are considered to be present in low abundance (<1%) by physical counts [11] and by sequence abundance [12–15]. Even at low abundance, metazoans have been associated with the global decline of microbialites [16, 17]. Metazoan grazing following the Cambrian explosion has been correlated to a drop in stromatolite diversity about ∼500 to 800 Mya [16, 17]. However, the metazoan grazing hypothesis is hindered by a lack of fossil evidence. In addition, protist grazers presumably far predated the Cambrian [18]. Another major problem is that unicellular or soft body grazers would leave no obvious fossils, resulting in a virtually nonexistent fossil record [18]. Additionally, despite their hypothesized role in the decline of microbialite systems, eukaryotes are vital to a healthy, functional microbialite community. A number of eukaryotic taxa have been shown to play a role in the accretion of microbialites. Photoautotrophic eukaryotes aid in the organosedimentary formation process of microbialites via photosynthesis-induced carbonate precipitation. Extracellular polymeric substances (EPS), heavily produced by protists such as diatoms, are very abundant in microbialites and are important for microbialite longevity by supporting sediment stability and resisting erosion [19]. Previous studies have documented diatoms from the Fragilarophyceae and Bacillariophyceae classes forming networks which can trap sediment grains and eventually constitute a stromatolite framework with the aid of cyanobacteria [20, 21]. Diatom-rich subspherical aggregates full of EPSs have also been described as a major contributor of carbonate precipitation in the hypersaline Laguna Negra (Argentina) microbial mats [22]. Eukaryotes such as Metazoans, Stramenopiles, Alveolata, Amoebozoa, Chlorophyta, and Rhodophyta were also found to be metabolically active in various microbialites across Highborne Cay and Shark Bay [23]. Another study looking at Lake Alchichica, a saline, alkaline crater lake in Mexico, identified green algae (predominantly a single Chlorophyte OTU) and diatoms as the primary protistan oxygenic photosynthesizers of microbialites. while also noting the presence of heterotrophic eukaryotes such as Fungi, Choanoflagellida, Nucleariida, Amoebozoa, Alveolata and Stramenopiles [12].

Notably, when characterizing the microbial communities of microbialites external factors such as location and salinity may drive changes in community composition, however the functional capabilities of these distinct communities remains mostly consistent [24]. Thus it is important to consider the context of the microbial interactions between the microbes when assessing the microbial community. Salinity can be a major driver of microbialite community composition and is also presumed to determine lithification rates in microbialite communities by inhibiting the growth of certain microbes [25, 26]. A study looking at the diversity of bacterial communities along a variable physicochemical gradient concluded that not only is salinity the main determinant of microbial community composition, but it also drives large scale changes in the metabolic potential of the bacteria [27]. A similar study of protist communities along a salinity gradient also found salinity to be the main driver of community composition [28]. By comparing the eukaryotic community assemblages of saline vs freshwater microbialite communities, we can uncover the underlying taxonomic similarities and differences in these functionally convergent communities.

Here we examined the eukaryotic diversity of freshwater microbialites (located in Kelly Lake and Pavilion Lake; Southwestern BC) in the context of other microbialites from the marine environment (Shark Bay, Australia and Highborne Cay, Bahamas) to conceptualize the emergence and decline of global microbialite communities as it relates to factors such as metazoan grazing and salinity. To this end, 18S rRNA (SSU) gene metabarcoding was performed to characterize the microbial communities of the freshwater sites. Both Kelly and Pavilion Lakes have microbialites that form across depths, to account for this we performed sampling across a depth gradient. These communities were analyzed in a meta-analysis incorporating the publically available 18S SSU gene barcode data from Highborne Cay and Shark Bay, which represent saline microbialite communities [23]. Our results highlight the similarities and differences within the metazoan and protist communities of freshwater vs saltwater microbialites providing further evidence for their importance in these ancient systems.

## Materials and Methods

### Site description and water chemistry

Pavilion Lake (50°51’ N, 121°44’ W) and Kelly Lake (55° 15′ 0″ N, 120° 2′ 0″ W) are mountain lakes in British Columbia, Canada, nestled in a steep-walled valley [29, 30]. Their major limnological characteristics have previously been described by Lim et al., (2009). Important features include their low phosphate concentrations (∼3 µg/L), relatively high CaCO_3_ concentrations (∼182 mg/L), relatively high pH (∼ 8.4), and clear waters with light penetrating to the bottom [30]. The microbialites consist primarily of calcite thrombolites, whose morphology varies strongly with depth (**Figure 1**): Shallow domes resembling “open lettuce-beds” are found at ∼10 m depth, intermediate domes resembling “cabbage heads” appear at ∼20 m, intermediate-deep domes with hollow conical “asparagus-like” outcroppings at 25 m, and deep domes possessing “artichoke leaflets” appear at and below 25 m [29, 31].

**Figure 1.**
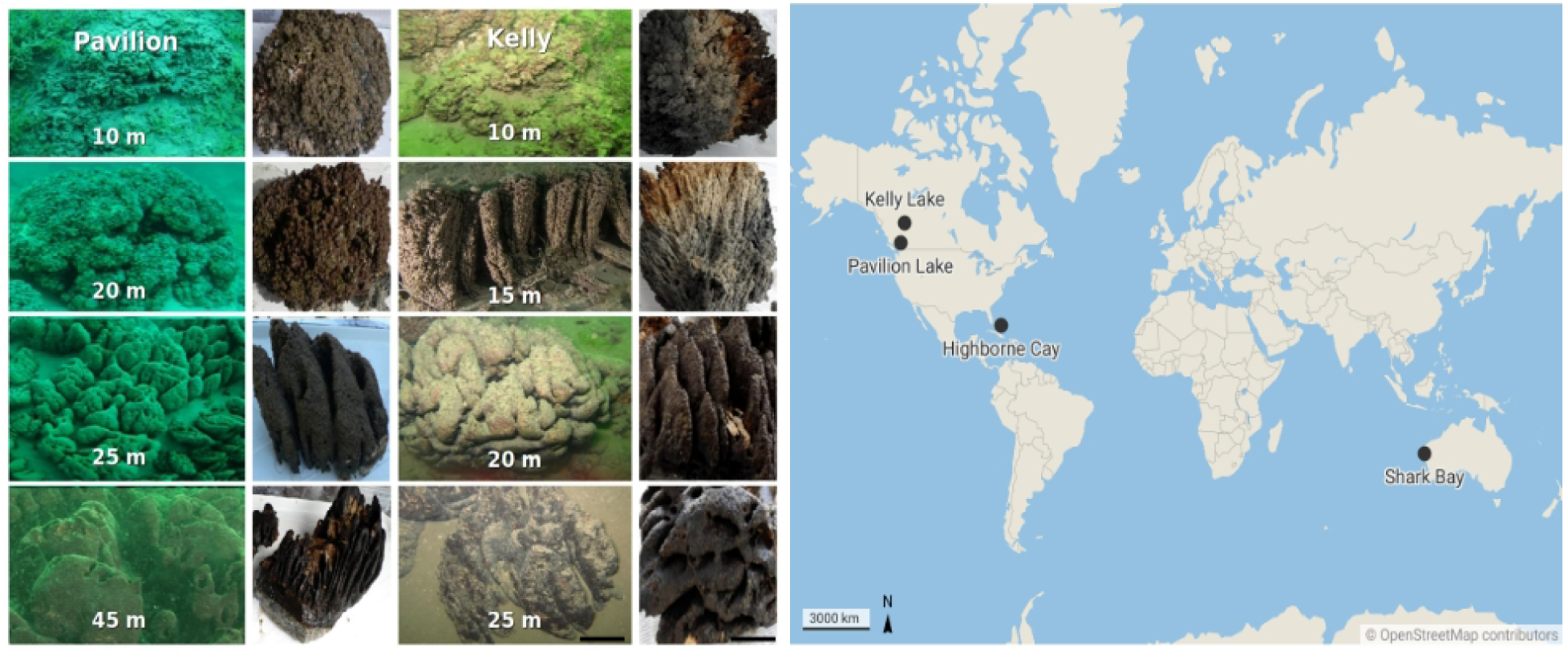
Sampling Map. Left) Microbialites of Pavilion and Kelly Lake structure by depth. Right) Map showing the locations of the four microbialite communities analyzed in this study.

### Sampling, DNA extraction, purity and concentration measurements

Microbialite samples were obtained during the summer of 2010 and 2011. Representative microbialites (∼10 kg) were sampled by SCUBA divers along a transect [Pavilion Three Poles (TP) site: 10, 20, 25 m, and Kelly site: 10, 15, 20, 25 m]. A manned DeepWorker submersible sampled the deepest sample in Pavilion Lake (Deep mound site: 45 m; 2010) (Nuytco Research Ltd., North Vancouver, BC, Canada). Divers collected microbialites in autoclaved bags then brought to the surface for dissection for Pavilion and Kelly depths 10-25 m [15]. The microbiome of the microbialites, sampled using a sterile razor blade, was used to scrape off 3 to 10 mm (∼5 g) across the surface of morphologically similar microbialites collected at each depth [15]. DNA was extracted on-site using a PowerMax® Soil DNA Isolation Kit (Mobio, Carlsbad, CA, USA). DNA concentrations were determined using a Nanodrop-3300 fluorospectrometer (ThermoFisher, Nanodrop Wilmington, DE) with PicoGreen^®^ reagent according to the manufacturer’s instructions (Invitrogen, Carlsbad, CA). Purity of extracted DNA samples was determined by absorbance (260/280 and 260/230 ratios) using a Nanodrop-1000 (ThermoFisher, Nandrop Wilmington, DE). The DNA extraction triplicates were PCR (using EUK1A and 516R primers) separately then pooled for each microbialite depth prior to Illumina library construction.

### Library construction and sequencing

For Illumina library construction, using purified PCR products that were end-paired, A-tailed (Lucigen NxSeq DNA prep kit, Middleton, WI) and ligated to TruSeq adapters (IDT, Coralville, Iowa); small fragments were removed twice using magnetic beads (Beckman Coulter, Danvers, MA). The resulting libraries were pooled, and sequenced using 250 bp paired-end reads on an Illumina MiSeq® by UCLA GenoSeq Core (Los Angeles, CA).

### Eukaryote sequence analysis for Kelly and Pavilion lakes

Reads were quality checked using FastQC and adapters + primers were trimmed off using Cutadapt then processed using the DADA2 pipeline [32, 33]. Only forward reads were used for the analysis, as these were shown to have comparable taxonomic coverage to the primers used in the marine microbialite samples (**Supplemental: PR2 DATA**). Sequences were filtered by length and quality score, then Amplicon Sequence Variants (ASVs) were inferred using DADA2’s sample inference algorithm. ASV tables for each dataset were then merged into a consensus table. Chimeras were removed, then taxonomy was assigned using the PR2 18S rRNA database v4.14.0 [34]. Due to a high number of low-resolution taxonomically unresolved ASVs, a BLASTN search was performed using the PR2 database with bacteria and archaea sequences added. Any sequences matching to bacteria or archaea were then removed from the analysis. Any ASV with an e-value (<3 x 10^-170^) and sequence similarity of 97%^+^, 95-97%, and 90-95% were reassigned to the species, genus, and family level, respectively. The final ASV table was then manually inspected for contaminants and possibly misassigned taxa. The final filtered dataset contained 2488 ASVs across the sampling sites. Alpha diversity plots were generated using the microbiome R package (http://microbiome.github.com/microbiome). Relative abundance plots were created showing only the taxa with a median relative abundance above 0.01%. Beta-diversity between the communities were assessed using Aitchison distances and ANOSIM as these methods account for the compositional nature of microbiome data [35]. A weighted UniFrac was constructed for further community analysis across locations using a phylogenetic tree of ASVs constructed using FastTree [36]. Analysis of Compositions of Microbiomes with Bias Correction (ANCOM-BC) was used to find significantly enriched taxa between the freshwater and saltwater microbialites [37].

### Comparison to Shark Bay and Highborne Cay

18S rRNA gene pyrosequencing from 9 marine microbialites in Highborne Cay (HC), Bahamas and 6 samples from Shark Bay (SB), Australia [23]. Short Read Archive (SRA) accession numbers are SRA061992 (Highborne Cay, Bahamas) and SRA061825 (Hamelin pool, Shark Bay, Australia). We completed the same quality filtering and chimera detection as we did with the Pavilion and Kelly lake 18S SSU data. These ASVs were then merged with our other dataset in DADA2 before taxonomic assignment and further processing as described above for the Kelly Lake and Pavilion Lake samples.

### Data depositing

All sequence data used in this study is freely available and available for public access Open Science Framework (OSF, https://osf.io/g28ym/, Modern Microbialites Eukaryome). All code, scripts, data, and documentation used for the analysis of this study can be found on github (https://github.com/delCampoLab/microbialite_euks_2023).

## Results and discussion

An analysis of the protist communities (protists hereby referring to all ASVs not classified as metazoans or embryophytes) from each microbialite location showed that Highborne Cay had the most diverse and significantly different community compared to the other three according to Shannon-Weiner Diversity Indices. The other locations did not show a significant difference in Shannon-Weiner alpha diversity between locations, highlighting consistency in Shannon-Weiner alpha diversity across 3 fairly distant microbialite locations **(Figure 2A)**. Grouping the locations based on salinity revealed a significantly higher (p_adj_ = 0.005) Shannon-Weiner alpha diversity in the saltwater vs freshwater microbialites (**Figure 2B**). These results support the hypothesis that salinity plays a major role in structuring microbialite communities across space. Examining the beta diversity across the dataset, we found that both location (ANOSIM; 1000 permutations, R = 0.5278, p = 0.001) and salinity (ANOSIM; 1000 permutations, R = 0.669, p = 0.001) were major drivers of dissimilarity (**Figure 2C**). However, primer-bias may also be playing a role in the dissimilarities as it was also shown to be a significant driver of dissimilarity in our dataset (ANOSIM; 1000 permutations, R = 0.669, p = 0.001).

**Figure 2.**
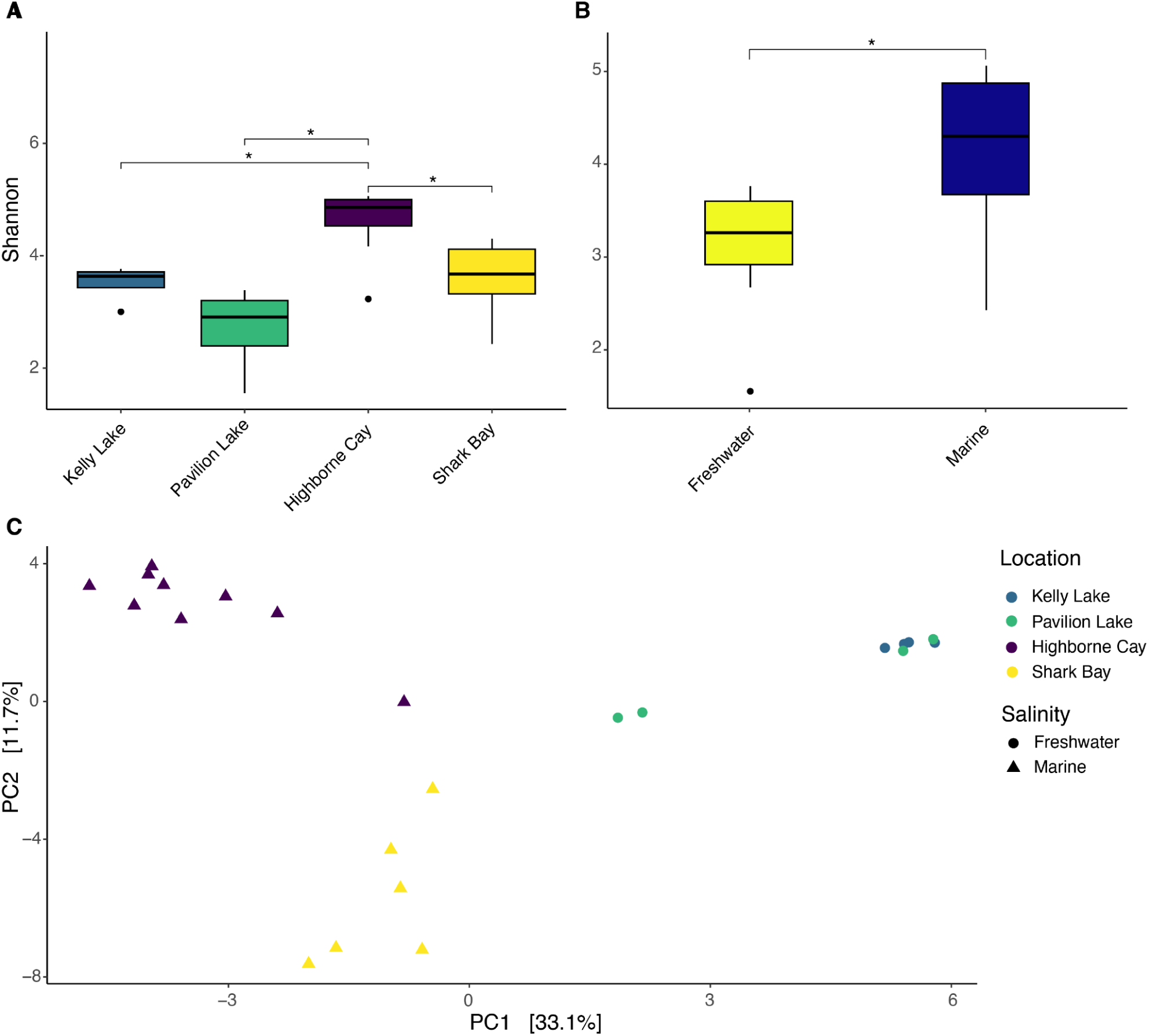
Alpha and beta diversities of microbialite protist communities. (A) Shannon-Weiner Alpha Diversity measurements for the four microbialites communities. (B) Shannon-Weiner Alpha Diversity measurements of the combined freshwater communities (Kelly & Pavilion Lake) and the combined marine communities (Highborne Cay & Shark Bay) showing a significant (P = 0.0053) difference in community diversity. (C) Aitchison Distance PCA of the microbialite communities included in this study.

Comparing the compositional makeup of the microbialite communities in this study, we see stark differences between the freshwater and marine locations. The microbialite protist communities at the freshwater locations were consistently dominated by Phaeophyceae, Ulvales-relatives, and Cladophorales which represented a contrast to the saltwater microbialite protist communities from Highborne Cay and Shark Bay which were dominated primarily by Raphid-pennate and Araphid-pennate diatoms (**Figure 3A**). Diatoms are key players in microbialite accretion, so their high abundance in these saline environments can potentially be attributed to this role. The saline locations also showed a high abundance of Pinguiochrysidaceae and Labyrinthulaceae, which were nearly absent in the freshwater locations. These two protist taxa are known to exist predominantly in marine environments. Likewise, the freshwater locations both had Tokophryidae ciliates, which were completely absent in the saltwater locations. Tokophryidae ciliates have been described as epibionts on lake-dwelling crustaceans [38], so they may not be microbialite-specific associates, but rather associates of the crustaceans which live on the microbialites. The previous publication on Highborne Cay and Shark Bay did note the high relative abundance of diatoms present in the microbialite communities, however the varied abundance of each diatom clade was not clear [23]. A recent study on the saline Lake Alchichica microbialite community in Mexico also showed the high abundance of Chlorophytes and Ochrophyta within the microbialites [39]. Some taxa that exhibited a similar distribution across all our locations include: *Thoracosphaeraceae*, *Pseudokeronopsidae*, *Holostichidae*, and *Oxytrichidae*. A weighted UniFrac was constructed to compare the microbialite communities of the locations in a phylogenetic context and showed that freshwater microbialites from Kelly Lake and Pavilion Lake clustered distinctly away from the marine microbialites of Shark Bay and Highborne Cay, further supporting the notion of salinity as a driving factor in microbialite microbial diversity **(Figure 3B)**. As for the metazoans, the freshwater locations showed a strong dominance of arthropods within their microbialite communities, with Insecta, Ostracoda, and Maxillopoda making up most of the metazoan community composition **(Figure 3A)**. In contrast, the saltwater metazoan communities were dominated by Nematoda, specifically Chromadorea. The relative abundance of metazoan reads to protists reads did not significantly influence the distribution patterns present in each of the locations **(Figure 3A)**. Overall, the consistency of these microbial protist community composition patterns across distant locations in our dataset emphasizes the core microbial members which are consistently associated with microbialites belonging to fresh and saltwater, despite other environmental variables.

**Figure 3.**
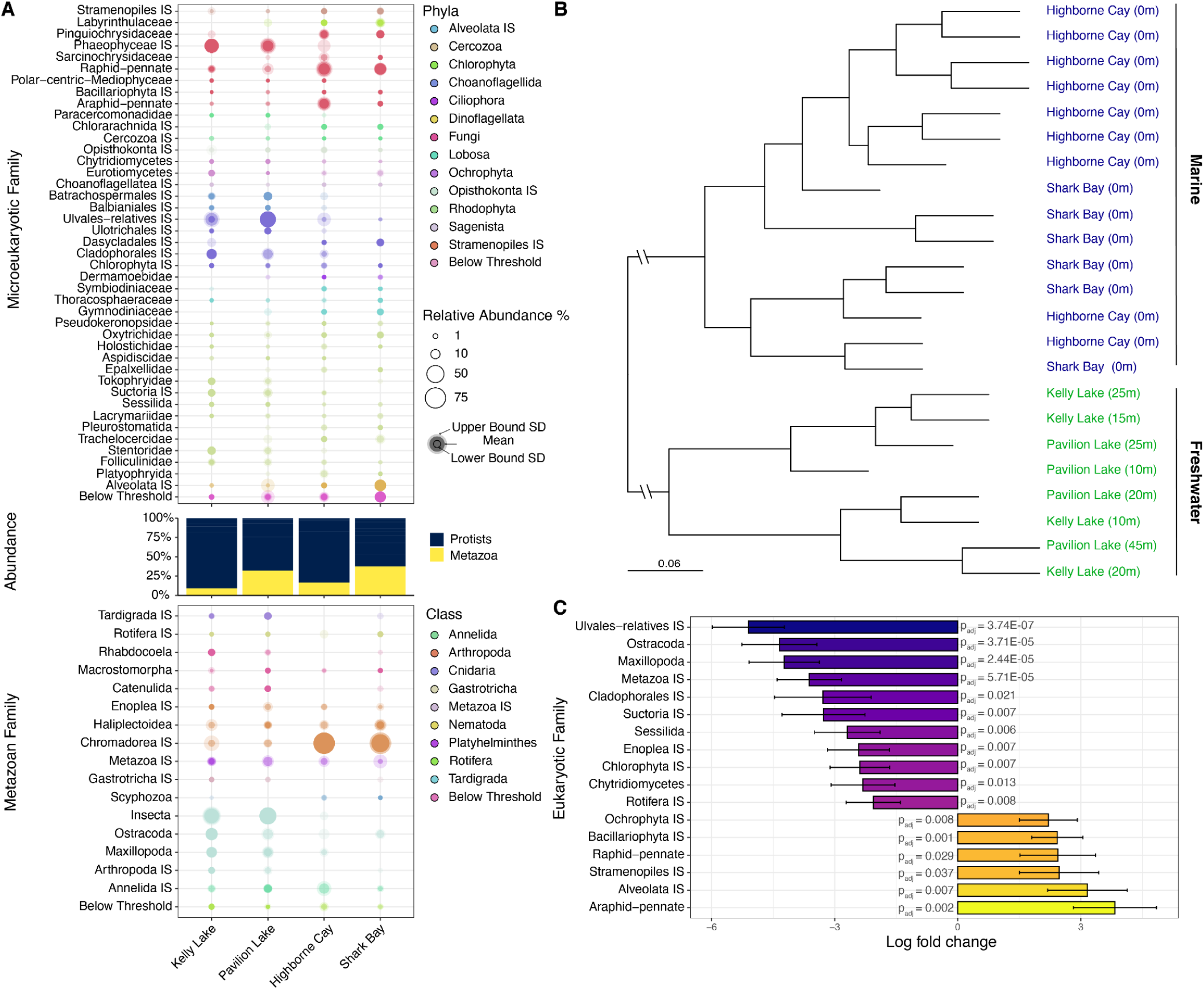
Protists and Metazoan microbialite communities. (A) Bubble plot depicting the relative abundance (mean ± SD) of eukaryotic communities of microbialites from the sampling locations. The proportion of protist to metazoan reads is also shown. (B) A weighted UniFrac of the eukaryotic communities of the samples used in this study, showing the clear delineation between marine (blue) and freshwater (green) microbialite communities. (C) Analysis of Compositions of Microbiomes with Bias Correction (ANCOM-BC) controlling for False Discovery Rates depicting the significantly (P < 0.05) differentially abundant eukaryotic taxa between marine vs freshwater microbialite communities.

Ulvales-relatives (Chlorophyta) were over ∼5 log folds more abundant in the freshwater locations than the marine microbialite locations (Figure 3C; p_adj_ = 3.74x10-7). Chlorophytes, specifically the filamentous green algae *Gongrosira*, along with cyanobacteria are associated with calcite precipitation in Pavilion Lake microbialites [40]. Interestingly, a recent phylogenetic analysis concluded that *Gongrosira fluminesis* which was previously identified as Chaetophorales actually belonged to Ulvales [41]. Carbonate-precipitating *Gongrosira* species have been found associated with freshwater microbialites around the world [40]. We hypothesize that our Ulvales-relatives detected in high abundance in Kelly Lake and Pavilion lake may play a significant role in microbialite carbonate deposition in freshwater microbialite communities around the globe along with cyanobacteria. Relatedly, chlorophytes were also recovered in high abundance from Lake Alchichica microbialites where they likely play a similar role [12]. Other ANCOM significant taxa in freshwater microbialites include Ostracoda, Maxillopoda, and Cladophorales, Sessilida, and Chytridiomycetes.

Marine microbialites were significantly dominated by diatoms, specifically Araphid-pennate (∼4 log folds more abundant; p_adj_ = 0.002) and Raphid-pennates (∼2 log folds more abundant; p_adj_ = 0.029). (**Figure 3C**). Again, the high abundance of diatoms in saline microbialites communities is consistent with previous observations and emphasizes their important role in microbialite stability through EPS production. Of note, the microbialite system in the saline and alkaline Lake Alchichica noted a significant presence of both chlorophytes and diatoms in the microbial communities [39]. These observations support the integral role these protists likely play in global microbialite communities.

To better understand freshwater microbialite protist communities, microbial community assemblages were assessed for both Kelly Lake and Pavilion Lake across a depth gradient revealing differing patterns of community composition between the two lakes. Phaeophytes are abundant at all depths in Kelly Lake, however Raphid-pennate diatoms tend to be more abundant at shallower depths (**Figure 4).** Phaeophytes, such as *Padina*, are capable of calcification [42], and thus may also contribute and complement the carbonate deposition role of Ulvales and cyanobacteria in freshwater microbialites. In Pavilion Lake this pattern is reversed with Phaeophytes being more abundant at deeper depths compared to shallower ones (**Figure 4**). As for Chlorophytes, Cladophorales show consistent assemblage patterns between the two lakes’ depth gradients being most abundant at deeper depths (**Figure 4).** Ulvales-relatives tended to be more abundant at shallower depths in both lakes as well (**Figures 4).** Groups of Cercozoans, Rhodophytes, and ciliates showed mostly consistent distribution across depths in both lakes. As for the Metazoan families along the depth gradient, Nematodes from the Haliplectoidea family and Chromadorea are more abundant in shallower microbialites than deeper ones in Pavilion Lake (**Figure 4).** Metazoan read abundance is lowest at 20 m depth in Kelly Lake (**Figure 4**), while Pavilion Lake shows a gradual decrease in metazoan read relative abundance as microbialites get deeper (**Figure 4).** While the two freshwater lakes share many environmental characteristics, Kelly Lake has much less microbial mat deposits and is generally considered a dead system in comparison to Pavilion Lake’s active microbialite communities.

**Figure 4.**
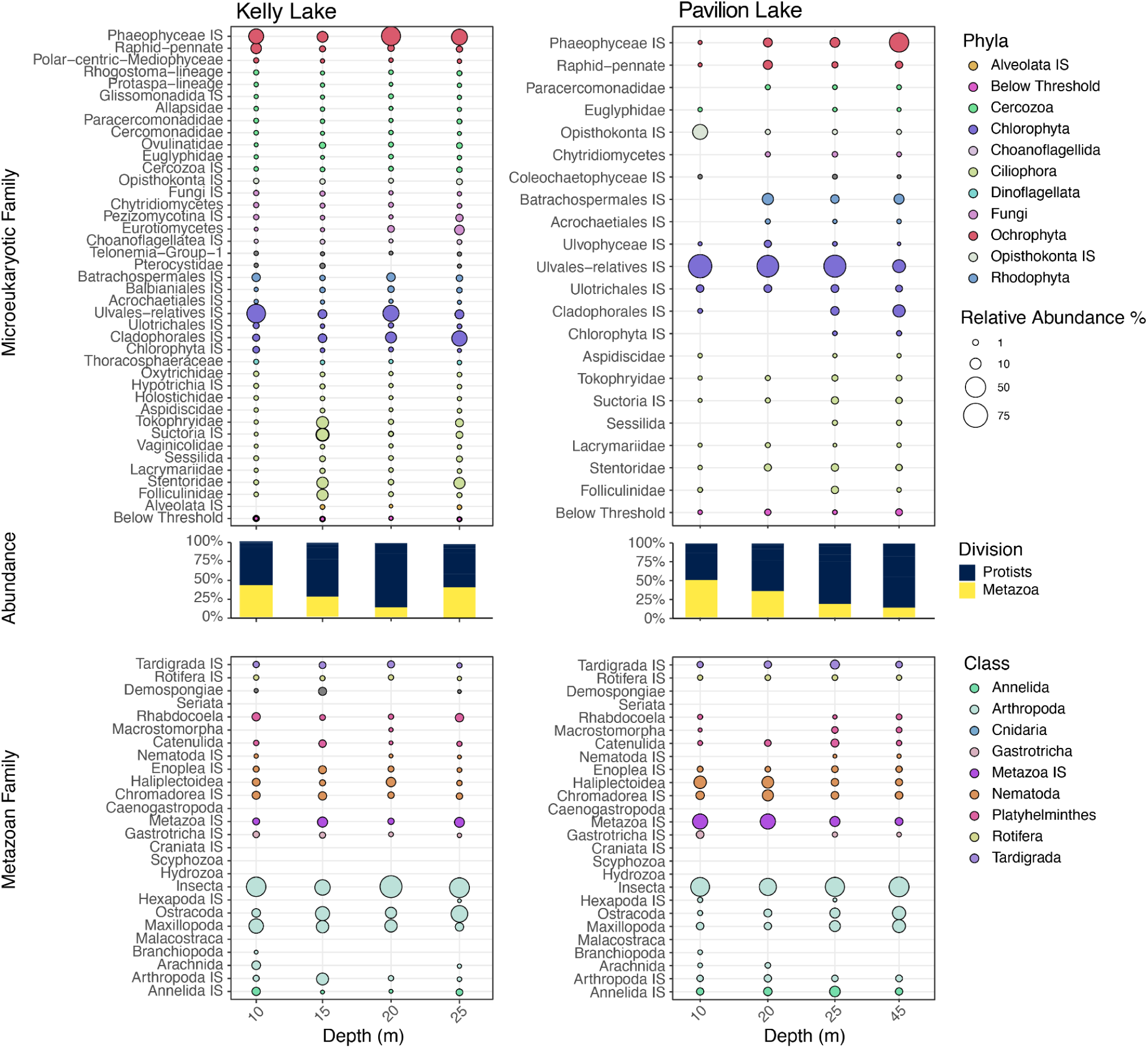
Depth profiles of protists and metazoan microbialite communities at Kelly & Pavilion Lake (>0.1%). The relative abundance of protist and metazoan orders along a depth gradient at Kelly Lake and Pavilion Lake, respectively. The proportion of protist to metazoan reads is also shown.

Generally, modern microbialites house a wide diversity of eukaryotes that are impacted by salinity on a global scale comparing marine to freshwater lakes. Chlorophytes green algae were significantly more abundant in the freshwater microbialites, while saltwater microbialites communities were primarily composed of pennate diatoms. Salinity appears to be a main driver of the eukaryotic community composition and structuring. Furthermore, more biogeochemical gradients should be evaluated for structuring the composition of these eukaryotes in modern microbialites.

## Acknowledgements

Here we provide many thanks to the entire Pavilion Lake Research Project (PLRP) research team, including Donnie Reid, Greg Slater, PLRP science divers, and the DeepWorker pilots for their help recovering samples from the lake. We also thank Curtis Suttle, Amy Chan, Danielle Winget, Kynan Suttle, Jan Finke, Marli Vlok, and Tyler Nelson for assistance with sample processing in the field. Microbialite morphology photographs in **Figure 1** are used with permission and copyrighted by Donnie Reid, Tyler Mackey, Amy M. Chan, Kynan Suttle, and Richard Allen White III. We are also grateful to Linda and Mickey Macri for hosting the PLRP and MARSLIFE projects and to the Ts’Kw’aylaxw First Nation and British Columbia Parks for their continued support of our research.

## Supplemental materials

**Supplemental Figure 1.**
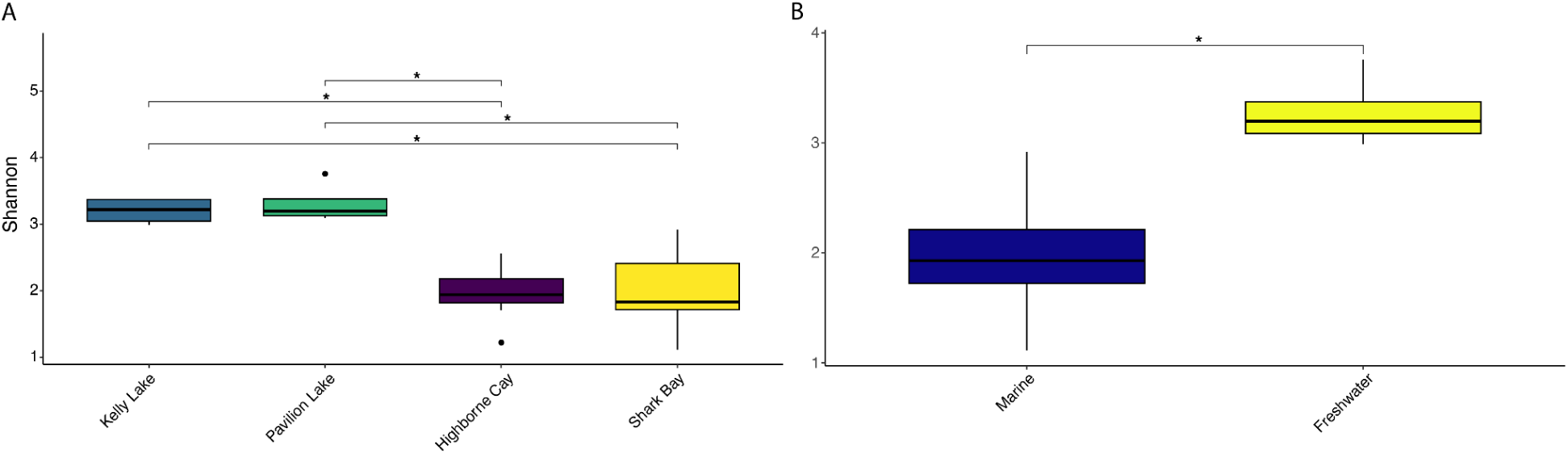
Alpha Diversity of Metazoan Communities by Location and Salinity. (A) Shannon-Weiner Alpha Diversity of the metazoan communities found across the sampling locations using only the metazoan-assigned reads. (B) Shannon-Weiner Alpha Diversity of the metazoan communities found in marine vs. freshwater biomes.

